# PDXNet Portal: Patient-Derived Xenograft model, data, workflow, and tool discovery

**DOI:** 10.1101/2021.10.15.464537

**Authors:** Soner Koc, Michael W. Lloyd, Jeffrey Grover, Sara Seepo, Sai Lakshmi Subramanian, Manisha Ray, Christian Frech, John DiGiovanna, Phillip Webster, Steven Neuhauser, Anuj Srivastava, Xing Yi Woo, Brian J. Sanderson, Brian White, Paul Lott, PDXNet Consortium, Yvonne A. Evrard, Tiffany A. Wallace, Jeffrey A. Moscow, James H. Doroshow, Nicholas Mitsiades, Salma Kaochar, Chong-xian Pan, Moon S. Chen, Luis Carvajal-Carmona, Alana L. Welm, Bryan E. Welm, Michael T. Lewis, Ramaswamy Govindan, Li Ding, Shunquang Li, Meenherd Herlyn, Mike Davies, Jack Roth, Funda Meric-Bernstam, Peter N. Robinson, Carol J. Bult, Brandi Davis-Dusenbery, Dennis A. Dean, Jeffrey H. Chuang, for the PDXNet Consortium Members

## Abstract

We created the PDX Network (PDXNet) Portal (https://portal.pdxnetwork.org/) to centralize access to the National Cancer Institute-funded PDXNet consortium resources (i.e., PDX models, sequencing data, treatment response data, and bioinformatics workflows), to facilitate collaboration among researchers, and to make resources easily available for research. The portal includes sections for resources, analysis results, metrics for PDXNet activities, data processing protocols, and training materials for processing PDX data.

The initial portal release highlights PDXNet model and data resources, including 334 new models across 33 cancer types. Tissue samples of these models were deposited in the NCI’s Patient-Derived Model Repository (PDMR) for public access. These models have 2,822 associated sequencing files from 873 samples across 307 patients, which are hosted on the Cancer Genomics Cloud powered by Seven Bridges and the NCI Cancer Data Service for long-term storage and access with dbGaP permissions. The portal also includes results from standardized analysis workflows on PDXNet sequencing files and PDMR data (2,594 samples from 463 patients across 78 disease types). These 15 analysis workflows for whole-exome and RNA-Seq data are freely available, robust, validated, and standardized.

The model and data lists will grow substantially over the next two years and will be continuously updated as new data are available. PDXNet models support multi-agent treatment studies, determination of sensitivity and resistance mechanisms, and preclinical trials. The PDXNet portal is a centralized location for these data and resources, which we expect to be of significant utility for the cancer research community.

## Introduction

Patient-Derived Xenograft (PDX) models are cancer models that support personalized medicine research and preclinical and co-clinical trials^1–5^. Specific PDX research areas include the study of sensitivity and resistance mechanisms, evaluation of new treatment options, and the study of tumor heterogeneity. The PDX research community is rapidly growing, with PDX-generated data being the preferred support for proposing human clinical trials^6^. In 2017, the National Cancer Institute (NCI) funded the PDX Development and Trial Centers (PDTC) research network (PDXNet, <u>pdxnetwork.org</u>) consortium to accelerate PDX research by developing new PDX models across cancer types, identifying new multi-agent treatments to bring forward into clinical trials, generating complementary RNA-Seq and whole-exome sequencing data, and increasing the ethnic diversity of publicly available PDX models.

PDXNet was also charged with developing collaborative research projects involving the 6 different PDTCs to advance PDX science. Each of the PDTCs came into PDXNet with its own home-grown data standards, data analytic pipelines and workflows. To facilitate collaboration, the disparate processes and databases required harmonization at many different steps, so that data from centers could be combined and analyzed efficiently. The harmonization goal was achieved through the creation of the PDXNet portal and the analytical tools within it. The PDXNet portal resources created by this effort enabled the successful completion of several collaborative research projects ^7–10^ and are supporting many others.

In addition to facilitating PDXNet research, an additional benefit of the PDXNet Portal is to make the PDXNet-generated data and workflows of the PDXNet Portal available as a public resource. These data will support cancer research by increasing the quantity and diversity of PDX data available and decreasing the effort required to analyze PDX sequencing data. We present the PDXNet Portal as a utility for PDXNet data for the larger scientific community.

There are several existing public PDX resources that complement the PDXNet Portal. Launched in 2012, the NCI Patient-Derived Model Patient Repository (PDMR)^11^ collects and develops PDX models and associated standardized sequencing data (RNA-Seq and whole-exome), with the goal of supporting academic and industry research. The PDMR maintains a publicly available database of models and an File Transfer Protocol (FTP) site for accessing sequencing data. Another resource is PDXFinder^12^. PDXFinder is an online resource that aims to harmonize internationally generated PDX models and their associated metadata. PDXFinder is a collaboration between the European Molecular Biology Laboratory’s European Bioinformatics Institute (EMBL-EBI) and the Jackson Laboratory. A key component of data harmonization in PDXFinder is the PDX minimal information standard (PDX-MI)^13^, which allows for standardized PDX information exchange. PDXFinder employs PDX-MI to support complex model searches that enable users to identify model descriptions and subsequently link to model information and a request form. EuroPDX is a consortium of eighteen non-profit cancer institutes that collaborate and coordinate PDX model development and access to improve cancer patients’ standard of care. Next, the EuroPDX Data Portal (https://dataportal.europdx.eu/) is a resource that provides information about PDX models generated by EuroPDX researchers and clinicians. Lastly, the Baylor College of Medicine PDX Portal (https://pdxportal.research.bcm.edu/) provides access to breast cancer, leukemia, pediatric liver cancer, pancreatic cancer, and sarcoma model collections.

The primary aim of the PDXNet Portal is to support the Cancer Moonshot^14^ model and data sharing goals^15^. The PDXNet Portal facilitates the distribution of resources, complementary data analyses, and developed tools. The resources generated include data collections and standardized bioinformatics workflows. Complementary data analyses such as data quality control analyses that support data-use are also available from the portal. Tools developed to support the use of the data (e.g., workflow cost estimation) are also accessible. Integration with the NCI Cloud Resource, the Cancer Genomics Cloud powered by Seven Bridges (CGC) ^16^, allows approved researchers to directly analyze PDXNet data or use developed workflows on private data. This manuscript details the PDXNet portal features that provide a gateway for identifying and accessing resources generated by the PDXNet community.

## Portal Design and Organization

The PDXNet Portal is designed to support coordination with other resources, including the PDMR, the CGC, and the NCI Cancer Data Service (CDS)^17^. Currently, the PDXNet portal references PDMR model information, genomic, transcriptomic, and tumor volume response data used in PDXNet research activities. The CGC serves as a PDXNet data staging area supporting data harmonization, standardized data processing, and research activities. The PDXNet Portal supports submission of sequencing data to the CDS to provide long-term research access to PDXNet data resources. The CDS is part of the NCI Cancer Research Data Commons which aims to store data resources generated by NCI-funded research. The CDS is available from across the NCI data infrastructure through a dbGaP access mechanism. The PDXNet portal augments the dbGaP submission process, through an administrative feature for generating data reports and through a dbGaP submission tool written to support CDS submissions. The PDXNet Portal aims to use data standards when they exist, supporting both the PDX-MI standard^13^ and the PDMR data structures^11^. These existing data structures allow for collaboration and information sharing with existing PDX resources.

## Portal Features

The PDXNet Portal is a publicly accessible website (https://portal.pdxnetwork.org/) with the primary function of providing access to the PDXNet models and information on how to obtain sequencing data. We extended the portal’s primary mission to include additional resources, including supporting access to the PDMR sequencing file data set, a PDXNet hematoxylin and eosin stain image data set, and PDMR tumor volume data. Below, we describe the features and sections of the Portal in detail.

### PDXNet Portal Landing Page

The PDXNet Portal Landing page includes an overview and summary panel of contents (Figure 1). The Portal overview identifies the primary PDXNet funding sources and participants. In summary, the NCI Cancer Therapy Evaluation Program (CTEP) funds four PDTCs and the PDCCC, whereas the NCI Center to Reduce Cancer Health Disparities funds two PDTCs (see Table 1 for additional details). The portal directs questions and requests for additional information to the PDXNet website at https://pdxnetwork.org.

**Table 1.** PDXNet Development and Trial Centers (PDTC) and the PDX Data Commons and Coordinating Center (PDCCC)

| PDXNet |  |
| --- | --- |
| PDX Development and Trials Centers (PDTC) |  |
| HCI-BCM* | Huntsman Cancer Institute and Baylor College of Medicine |
| MDACC | MD Anderson Cancer Center |
| WUSTL* | Washington University at St. Louis |
| WISTAR* | Wistar Institute and MD Anderson Cancer Center |
| BCM<sup>&</sup> | Baylor College of Medicine |
| UCDAVIS<sup>&</sup> | The University of California at Davis |
| PDX Data Commons and Coordinating Center |  |
| JAX-SB*<sup>&</sup> | Jackson Laboratory and Seven Bridges |

\* NCI Cancer Therapy Evaluation Program Funding
<sup>&</sup> NCI Center to Reduce Cancer Health Disparities Funding

**Figure 1.**
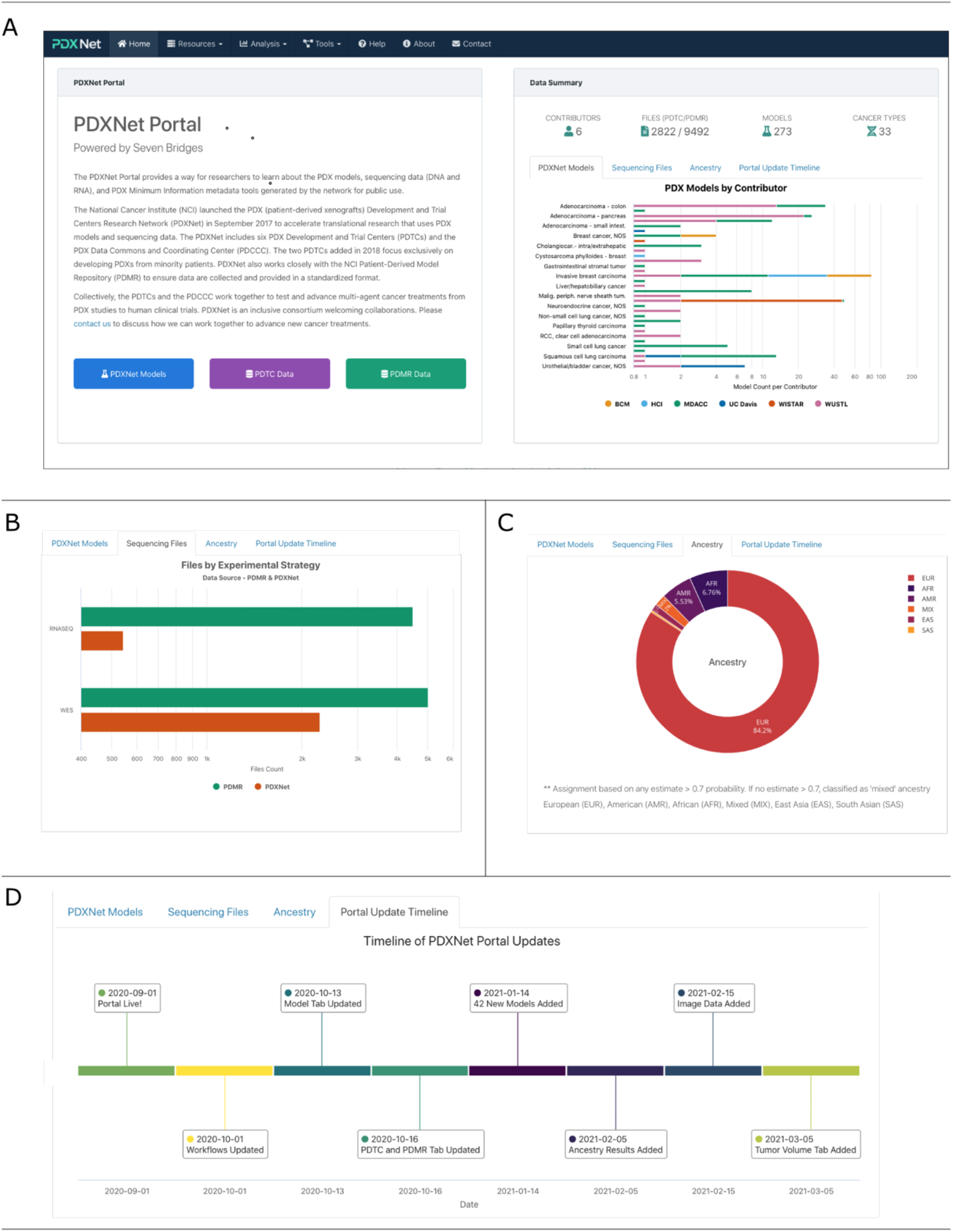
PDXNet Portal Landing Page. Views from the PDXNet Portal Landing Page. **(A)** The initial PDXNet Portal Landing Page. **(B)** Experimental strategies (whole-exome and RNA-Seq) for the PDXNet (green) and PDMR (red) sequencing datasets. **(C)** Computed ancestry in a pie chart. Ancestry is classified in the following categories: European (EUR-Red), African (AFR-Blue), American (AMR-Purple), Mixed (MIX-Orange), East Asian (EAS-Light Purple), South Asian (SAS-Yellow). **(D)** Major portal updates in a timeline starting in September of 2019 through March 2021.

The data summary panel on the right side of the screen allows the reader to review model and data summaries and the portal update timeline. The data summary panel lists the number of PDTCs contributing data, the number of files uploaded by the PDTCs and available from the PDMR, the total number of models, and the number of cancer types represented in those models. Tabs allow the reader to review summary figures for PDX Models by cancer type and contributing PDTC, sequencing files by experimental strategy, ancestry, and the Portal Update Timeline.

### Resources

The PDXNet Portal resources section includes pages that describe models, data (genomic, transcriptomic, and image), and analysis workflows made available on the CGC by the PDXNet consortium. The CGC based analysis workflows are a significant resource developed by the PDXNet community, allowing for reproducible and standardized analysis of PDX data^8^. The resource section also highlights data mirrored from the PDMR sequencing data repository to the CGC to support research activities. Lastly, we provide interactive plots and tables of sequencing data information for the PDXNet and PDMR sequencing data sets on the CGC. The PDXNet portal presents each resource (e.g., PDXNet, PDMR, workflows) on a separate page.

#### PDXNet Models

The PDXNet Portal Models tab summarizes verified model submissions to the PDMR made by each PDTC (Supplement Figure 1). The PDXNet models are a primary consortium deliverable. Each model submitted by a PDTC to the PDMR includes a completed model submission form that details the general PDX information, model-specific information, and tissue implantation details. Metadata are consistent with the PDX-MI and the PDMR data format. The metadata includes model id information to facilitate search and cross referencing to related PDMR models.

To date, PDXNet researchers have submitted 334 models to the PDMR across 33 cancer types. The most prevalent model cancer types include invasive breast carcinoma (30.8%, 103), melanoma (20.1%, 67), adenocarcinoma – colon (12.3%, 41), and adenocarcinoma – pancreas (7.8%, 26). See Table 2 for additional details.

**Table 2.**
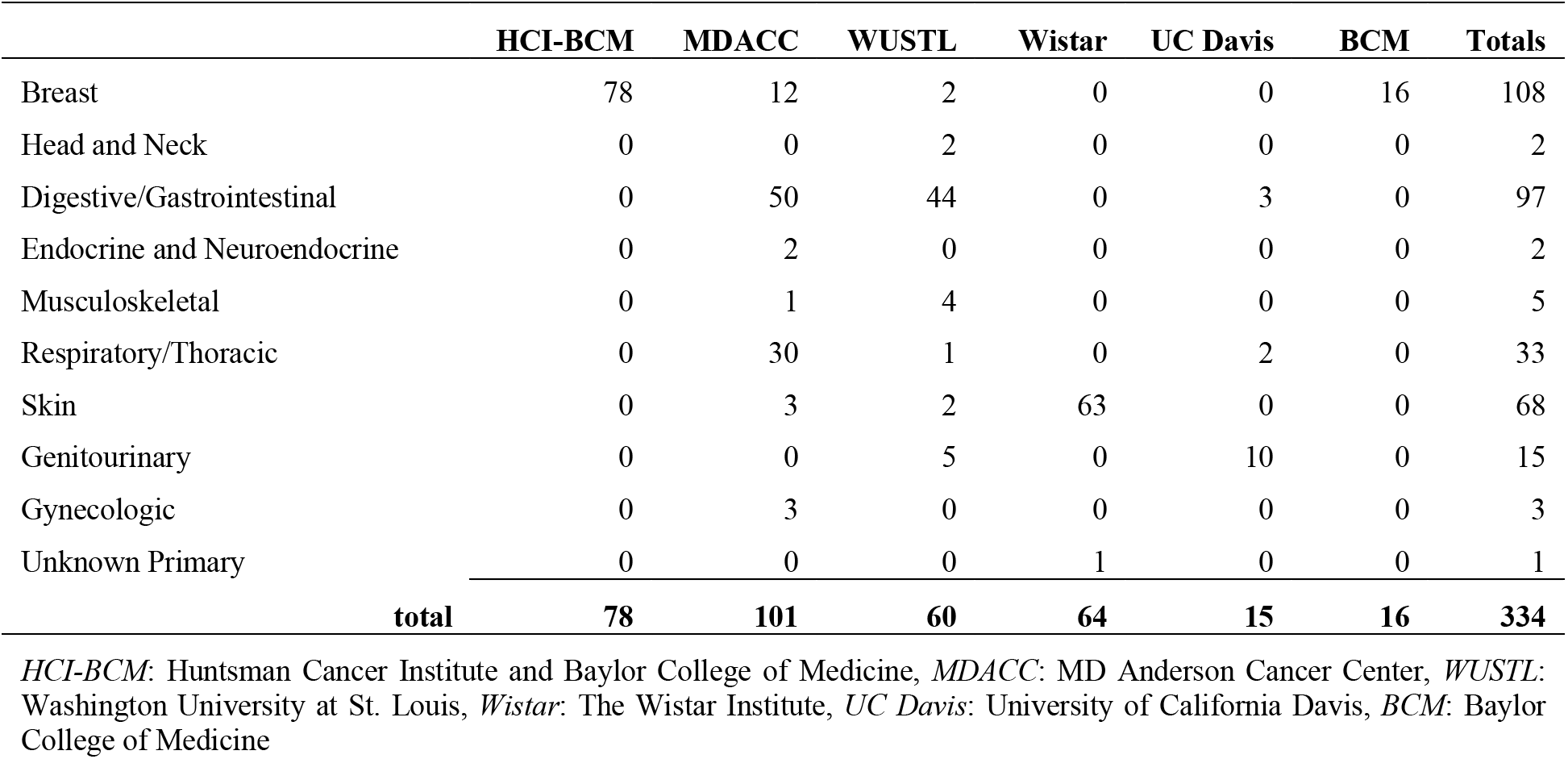
PDX models generated by PDX Development and Trials Centers.

#### PDTC Sequencing Data

The PDXNet Portal summarizes sequencing data submitted by the PDTCs for intraconsortium sharing and for public sharing (Figure 2). The PDXNet Data Collection - PDTC tab presents the core PDXNet sequencing data set. We processed submitted sequencing data with standardized workflows (e.g., whole exome capture; additional) according to a written standard operating procedure provided on the CGC. See the workflow section for description of the workflows used for standardized processing.

**Figure 2.**
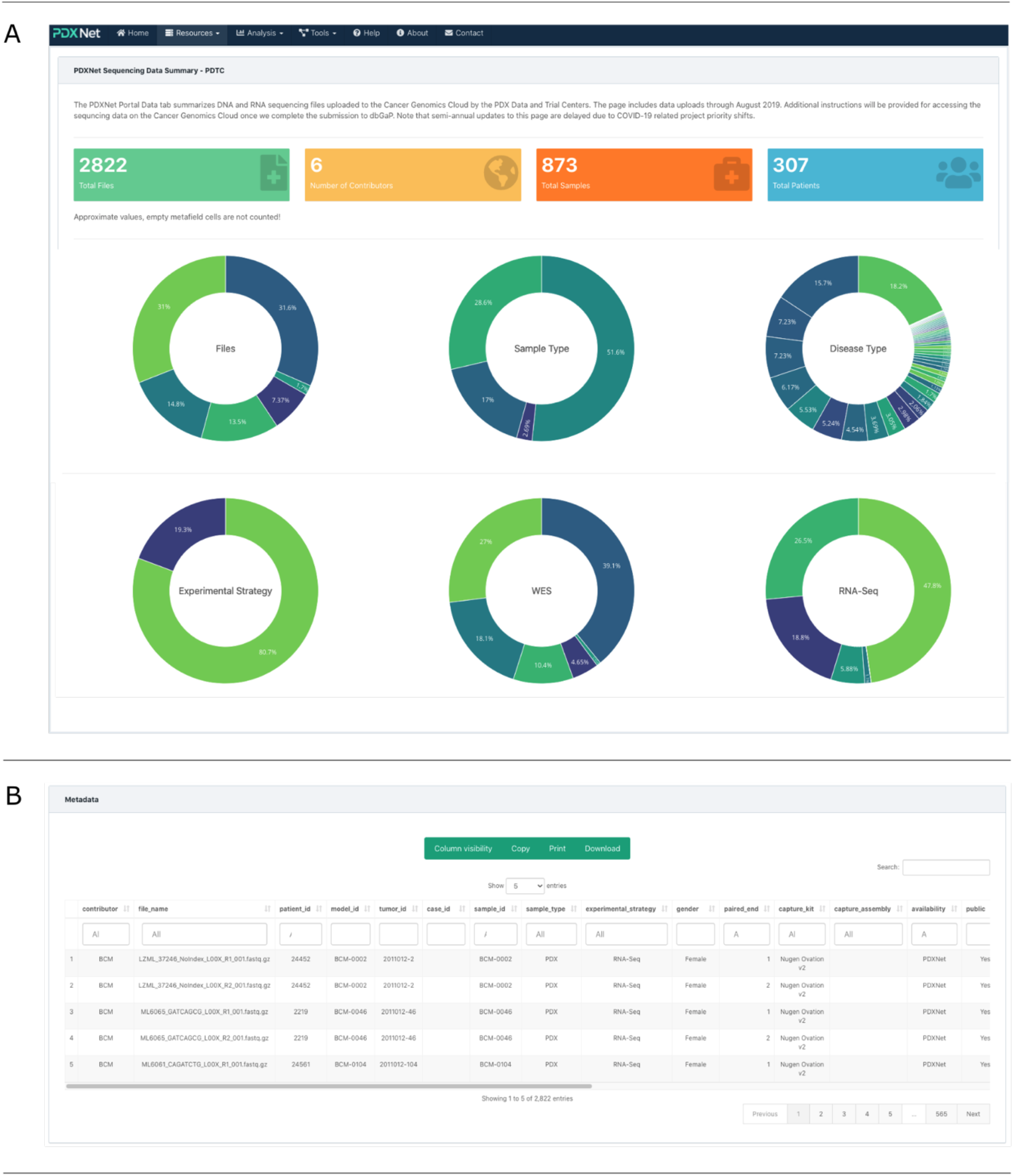
PDXNet sequencing data page on the PDXNet Portal. Components of the PDXNet sequencing data page. (A) Panel shows summary statistics including number of sequencing files (green), contributors (yellow), total samples (orange), and total patients (blue). Also, shown are donut plots for contributors, sample types, disease type, experimental strategy, WES contributors, and RNA-Seq contributors. (B) Panel shows metadata for the PDXNet sequencing data in a spreadsheet format. The interface supports searching and sorting metadata. Users can copy, print, and download metadata into accessible formats.

PDXNet researchers contributed 2,822 total sequencing files that include both whole-exome (80.7%, 2,278) and RNA-Seq (19.3%, 544) data. Six institutional contributors submitted 873 samples from 307 patients. The sequencing sample types include PDX (51.6%, 1457), tumor (28.6, 808), normal (17%, 480), and blood (2.7%, 76). The most prevalent diseases represented among the samples include breast (42.1%, 1,189), lung (12.8%, 362), pancreas (6.2%, 174), colon (5.8%, 164). Metadata provided by centers did not include disease information for 18.2% (515) samples (see Table 3 for additional information).

**Table 3.**
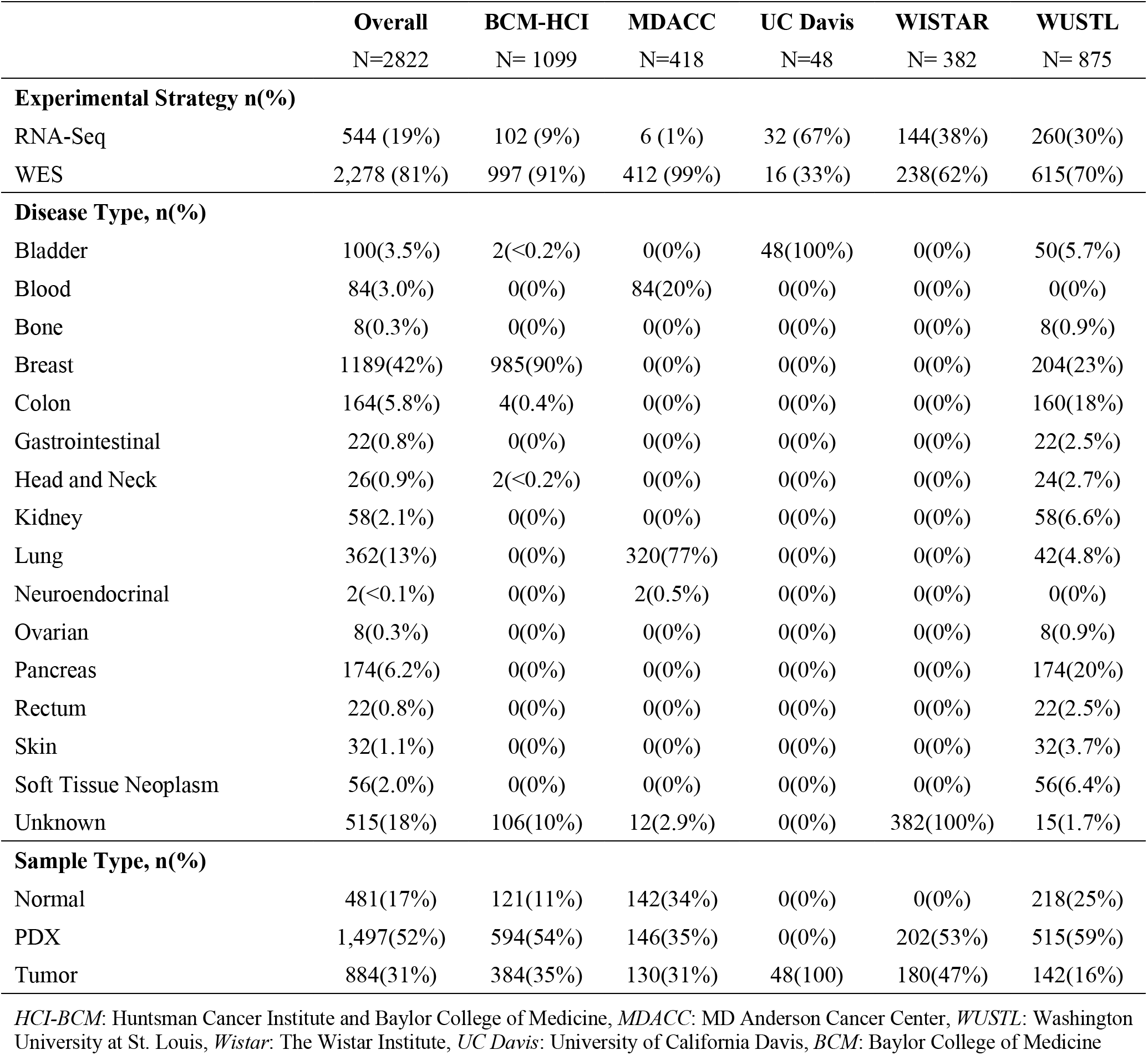
Sequencing data files generated by PDX Development and Trial Centers.

#### PDMR Sequencing Data

The PDXNet Portal Model tab summarizes sequencing data transferred from the PDMR FTP server to the CGC as of August 2020 (Supplement Figure 2). The PDMR generates whole-exome and transcriptome sequencing data from models submitted according to tissue collection best practices and model quality control practices^11^. Molecular characterizations include whole-exome sequencing and gene expression profiling. We processed the PDMR sequencing data with standardized workflows as for the PDTC data.

The PDMR sequencing dataset on the CGC includes 9,492 paired-end sequencing files that include both whole-exome (52.8%, 5,012) and RNA-Seq (47.2%, 4,480) data (See Supplement Table 4). The data set includes 2,594 samples from 445 patients covering 34 disease types. The sequencing sample types include PDX (82.7%, 7,846), primary tumor (5.7%, 542), normal germline (5.5%, 520), and organoid (3.2%, 304). The most prevalent diseases represented among the samples include colon (21.1%, 2,002), head and neck (11.6%, 1,102), soft tissue neoplasm (10.1%, 958), skin (8.7%, 828). Due to the size and cost associated with data transfers, synchronization between the PDMR sequencing database and the CGC dataset is done periodically. The PDMR data webpage has the most updated list of available PDMR sequencing data processed with standardized PDXNet workflows.

#### PDMR Image data

The PDXNet Portal Image tab summarizes hematoxylin-eosin stain (H&E) image data provided by the PDMR (Supplement Figure 3). The PDMR image data on the CGC includes 593 images scanned from PDX (93.8%, 556) and primary tumors (6.2%, 37). The images correspond to 593 samples taken from 92 patients across 37 disease types. The PDX passages ranged from P0 to P6 with the top four passages corresponding to P1(41.4%, 225), P2 (24.1%, 131), P0 (22.8%, 124), and P3 (8.3%, 45). The PDXNet Portal currently supports 43 metadata fields that data submitters can populate upon submission (See Supplement Table 1 for the complete list).

#### Interactive Data Explorers

The PDXNet Portal data explorer allows users to interactively create summary tables from the PDXNet and PDMR sequencing datasets (Supplement Figure 4). The interactive table supports 10 table and chart types including simple tables, bar charts, line charts, and heat maps (see Supplement Table 1 for full list). Interactive tables also support 22 data summary options including count, sum, average, and variance (see Supplement Table 2 for the full list). The user drags and drops from 20 metadata fields to the table type area to construct the table. Metadata field examples include contributor, sample type, experimental strategy, and passage (see Supplement Table 3 for complete list).

#### PDXNet Workflows

The PDXNet Portal Workflows tab summarizes analysis workflows developed by the PDXNet community (Supplement Figure 5). We selected workflows for standard consortium-wide data processing and public release from those submitted by each PDTC after benchmarking with simulated and experimentally derived PDX data^8^. Since the initial public release, we have restructured the workflows to efficiently process normal (tissue), tumor-only, and tumor-normal data. These workflows are implemented on the CGC using the Common Workflow Language (CWL)^18^ with Docker containerized tools, which allows for easy sharing and analysis reproducibility. The PDXNet consortium developed a set of 15 workflows validated for processing of both whole-exome and RNA-Seq data (Supplement Table 4). We are sharing these validated and tested workflows with the broader community via the CGC Public Apps Gallery (https://cgc.sbgenomics.com/public/apps#q?search=pdx). The use of CWL allows these workflows to be portable to any CWL-compliant execution environment. The workflows collectively facilitate the analysis of whole-exome or RNA-Seq data via mouse read disambiguation, read alignment, variant calling or transcript quantification, and sample and cohort level quality control. For whole-exome data, we also compute copy number variation (CNV), tumor mutational burden (TMB), microsatellite instability (MSI), and homologous recombination deficiency (HRD) during standardized processing. A full explanation of inputs, outputs, and data processing steps for each workflow is provided on the CGC in the respective description section.

### Analysis

The PDXNet Portal analysis section includes several metrics derived from primary data sources and are described below in more detail. These results were generated from standard processing analysis workflows or through PDXNet research activities^8^, and we provide these analyses to support independent research by the broader research community.

#### Ancestry Analysis

The PDXNet Portal Ancestry Analysis page summarizes genetic ancestry analysis for datasets on the portal (Supplement Figure 6). We compute ancestry with SNPweight^19^ using a reference dataset generated from the 1000 Genomes Project Phase III^20^. We classify each sample into one of five categories, which correspond roughly to the concept of “continental ancestry.”^21^ These categories include European (EUR), African (AFR), American (AMR), East Asia (EAS), and South Asian (SAS). Samples that could not be confidently assigned to one of these categories are labeled Mixed (MIX). On the left side of the page, ancestry data filters allow the user to select the data contributors, ancestry, and disease type. Applying the selected filter to the data regenerates the two summary figures. The first summary figure is a bar chart that shows the ancestry distribution for the selected disease types. The second summary figure shows a pie chart with each slice corresponding to the ancestry types chosen. Supplement Table 6 shows the summary of ancestry estimation from PDX Models submitted to the PDMR.

#### HRD-TMB-MSI Analysis

The PDXNet Portal HRD-HSI-TMB analysis page allows the user to filter and summarize three computational metrics generated from whole-exome sequencing data by PDXNet standardized processing (Figure 3). The three computed metrics are Homologous Recombination Deficiency (HRD), Tumor Mutational Burden (TMB), and Microsatellite Instability (MSI). HRD is computed with ScarHRD^22^ for matched normal data. TMB is calculated as the number of coding mutations that meet all quality criteria per Mb of the genome. Quality criteria are assessed using coverage, allele frequency, mapping quality, and strand bias. Variants included in the calculation are somatic and non-polymorphic, and are defined in SnpEff^23^ as ‘high’ or ‘moderate’ functional impact. As only a portion of the genome was sequenced, genome coverage (Mb) is calculated from the input target coverage BED file. MSI is calculated with MANTIS^24^ for samples with matched normal data, and calculated with MSIsensor2^25^ for tumor-only samples. For each metric, users can set data filters for the visualizations. The data filters, on the left side of the page, allow the user to select data contributor, sample type, experimental strategy, and disease type. Applying the selected filter to the data generates a boxplot chart displaying the selected metrics for each disease type chosen (Figure 3).

**Figure 3.**
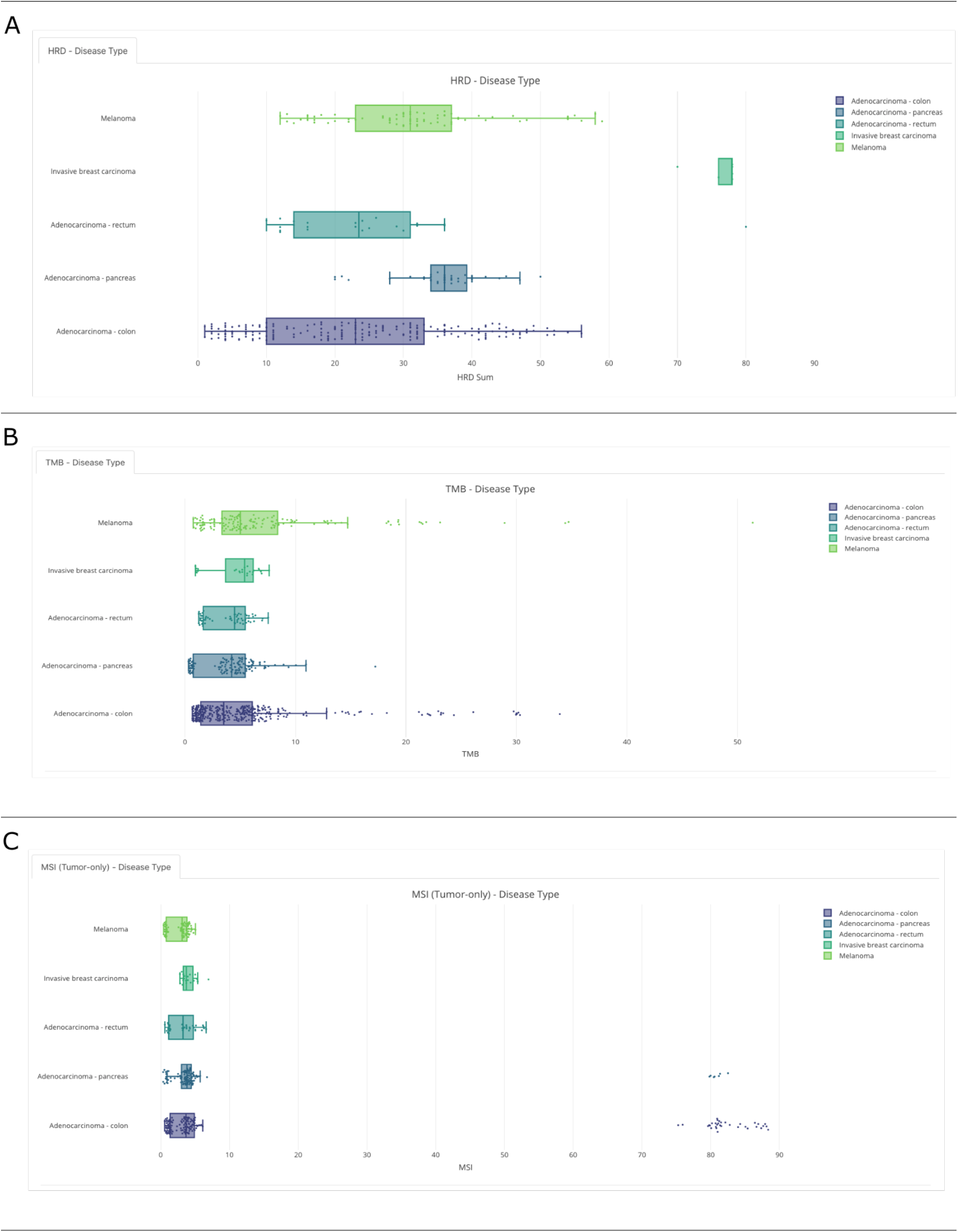
Examples figures generated from the HRD-TMB-MSI page on the PDXNet Portal. Plots generated on the PDXNet Portal HRD-TMB-MSI page **(A)** Plot of Homologous Recombination Deficiency (HRD) computed from sequencing data provided by PDXNet researchers. The plot shows HRD by disease type **(B)** Plot of Tumor Mutational Burden (TMB) computed from sequencing data provided by PDXNet researchers. The plot shows TMB by disease type. **(C)** Plot of TMB computed from sequencing data provided by PDXNet researchers, by disease type.

#### Tumor Volume Analysis

The PDXNet Portal Tumor Volume Analysis Page allows the user to visualize raw tumor volume growth data provided by the PDMR (Figure 4). The filters enable the user to select contributor, treatment, and disease type on the page’s left side. Applying the selected filter to the data regenerates the Tumor Volume and the Tumor Disease Types figure tabs. The currently available volume data is from 75 models representing 30 disease types, which were treated with seven possible agents (Supplement Table 7). The dataset has 17,920 volume measures from 89 treatment studies. The Tumor Volume tab allows the user to choose plot level (Animal and Treatment Arm) and plot pattern (multiple and combined), reorganizing the plots to correspond to select values. The Tumor Disease type tab plots a disease pie chart based on user selection (Figure 4).

**Figure 4.**
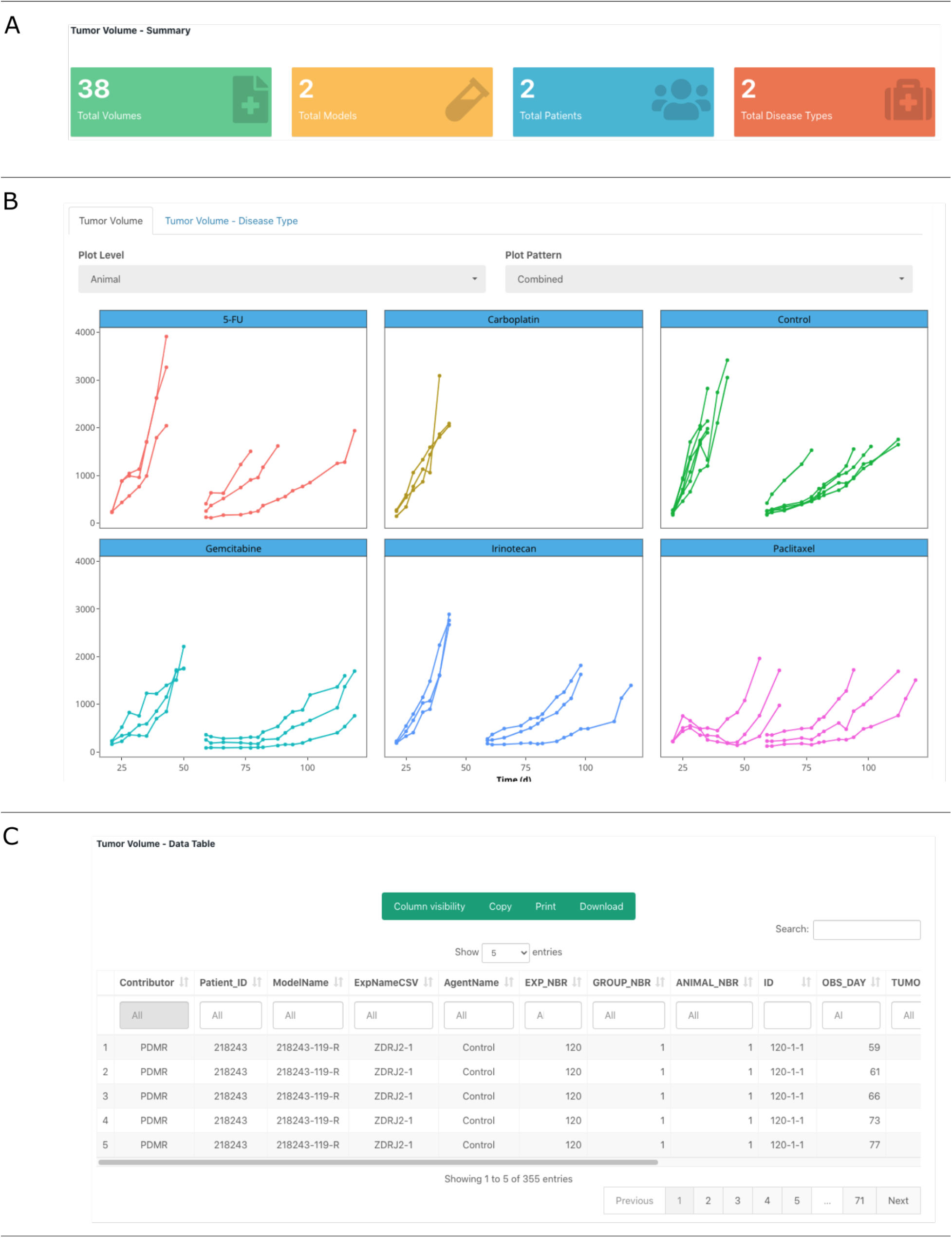
Tumor volume data page on the PDXNet Portal. Components of the PDXNet tumor volume page. The figure shows a filtered dataset. **(A)** Panel shows summary statistics including number tumor volume datasets (green), number of models in the selected dataset (yellow), total number of patients (blue), and total number of diseases represented (red). (B) Panel shows the tumor volume data organized by the treatment arm. The user can control plot level (animal or treatment arm) and plot pattern (multiple or combined)

#### Quality Control Analysis

The PDXNet Portal QC Analysis page provides plots and tabular results for selected QC metrics (Supplement Figure 7). The page displays QC metrics generated during the standardized data processing procedure for each relevant data type. The page provides sub-tabs showing whole-exome and RNA-Seq quality control metrics for a selected dataset PDXNet or PDMR. The whole-exome tabs plot mean target coverage, percent target bases with greater than 20% coverage, and percent duplication by data contributor. The RNA-Seq tabs plot percent usable bases, percent ribosomal bases, and percent correct strand reads. The plotted metrics, along with additional QC metrics are available in a table at the bottom of the page.

### Tools

The PDXNet Portal tools section includes several Portal specific tools developed to support present and future PDXNet and other independent general research activities.

#### Workflow Cost prediction

The Workflow cost prediction tool allows users to estimate the cost of processing their samples on the CGC with the PDXNet workflows. This tool uses prediction models (gradient boosting trees^26^) generated from 7,000 workflow runs. The user can select either whole-exome or RNA-Seq workflows and provide the number and optionally size of files to process. The calculator computes the storage and computation cost for processing the user defined dataset. The estimated costs assume the workflows were run on the CGC using spot instances. We expect the tool to allow users to estimate data storage and computational cost for their own analyses allowing for estimating grant budgets and budgeting lab expenses.

#### PDX Minimum Information Metadata – Creation

The generate metadata tab allows users to interactively generate a PDX minimum information metadata sheet (PDX-MI). As described above, the PDXFinder working group developed the PDX-MI as a standard for exchanging PDX information among institutions. The generate metadata tab allows the user to create a PDX-MI spreadsheet by stepping through data entry dialog boxes. The interface supports entry of patient information, treatment information, tumor information, model, and sequencing metadata. The user downloads the spreadsheet upon data entry completion, and no information is stored permanently on the PDXNet Portal site.

#### PDX Minimum Information Metadata – Validation

The validate metadata tab allows users to upload and validate a PDX-MI metadata spreadsheet. The user can review uploaded contents at the bottom of the page. The validation verifies that required metadata fields are present and that entries are valid. The validation feature generates a summary of required fields that includes percent completed and most common data entry per field. The validation feature reduces the amount of time necessary to review and check submitted PDX-MI spreadsheets.

### Implementation

The backend of the PDXNet Portal is an R-Shiny app hosted on a cloud-based server. The portal uses the PDMR and PDX-MI metadata standards. The PDMR and PDX-MI standards allow us to harmonize data across sources, quickly import data from the PDMR and other data sources, and exchange information with other PDX related portals. We also collect additional metadata required to facilitate computation on the CGC, including omics-related information. We use the Cancer Therapy Evaluation Program (CTEP) disease classification to standardize disease entries although we initially accepted institutionally-defined disease classification. In the cases where a standard does not exist; we collect sufficient metadata required to display and process the data source. For example, we take a minimalistic approach to managing tumor volume and H&E image data. Several PDXNet teams are working towards the development of best practices for these data types. Until these best practices are published, we will evolve these operational standards to support harmonization and analysis.

The PDXNet Portal team updates information on the portal semi-automatically using the same data model as the PDMR, allowing PDXNet to sync with PDMR model information. The PDMR provides regular updates to PDXNet on PDTC model submissions to update the PDX Model’s page. We receive sequencing data upload updates from the PDTCs and the PDMR, and we have developed scripts for extracting PDXNet standardized processing results, allowing for quality control information and computational metrics to be tabulated for semi-automated PDXNet Portal updates. The PDXNet Portal source code will not be made publicly available for security reasons. Future PDXNet Portal versions will support controlled access sign-in to provide links to controlled files.

### Data Availability

Each PDXNet Portal data tab allows users to download metadata. Data for smaller data types such as tumor volume data and computed metrics (ex. HRD and TMB) can be downloaded directly from the portal. For larger data types, please request data from the PDXNet Portal’s Contact page. We will coordinate with PDTCs to make data available either directly or through dbGaP as required by the PDXNet data sharing agreement.

## Discussion

The PDXNet Portal is a vital component of the PDXNet consortium. The portal establishes a mechanism for public discovery of consortium-generated resources including models, data, and workflows. The portal allows researchers to examine the data, models, and metadata using integrated query features. These portal capabilities facilitate cancer data discovery, a core goal of the NCI Cancer Moonshot program. Additionally, the PDXNet Portal allows the consortium to manage data analysis projects by clarifying which data are available, their source, their quality, and their suitability for research projects and scientific questions. Within the PDXNet consortium, the PDXNet Portal functions as a centralized source of information for the status of the available models. The standardized sequence processing, quality control, and computation of common metrics (e.g., MSI, TMB, HRD, and genetic ancestry) further enhance data analysis, model use, prioritization of future models and data collection.

Enhancement of collaboration between researchers is a main objective of the PDXNet Portal. To accomplish this, we are integrating the PDXNet Portal and PDXNet data with existing NCI, NIH, and NCBI infrastructure. All data visible on the PDXNet Portal will also be available through the Cancer Data Service (CDS)^17^ through dbGaP^27^ access. Accessing PDXNet data on the Cancer Genomics Cloud^28^ allows users to perform sophisticated analyses utilizing cloud computing within an integrated bioinformatics ecosystem. By co-locating data and analysis, as well as integrating data management, this infrastructure can decrease the time required for researchers to perform analyses.

For large consortia such as PDXNet, metadata and secondary data types are often just as important as sequencing data for supporting impactful research. Examples of these additional data types include high-resolution images and tumor volume/drug response data. These data extend the types of problems researchers can address. We expect that the portal’s image and tumor volume functionality will expand as these datasets grow. Future iterations of the portal will include interactive exploration across data types allowing users to address complex research problems.

PDX models are widely used in cancer research, but there remain challenges in standards for data submission, access, and quality. The PDXNet Portal reflects PDXNet activities to implement such standards not only for sequencing data but also metadata and secondary data types. Another consideration requiring careful implementation is to balance data security versus ease of use. The PDXNet Portal will grow with new data and features as the PDXNet consortium continues to generate new models. Consequently, standardized processing and batch effects are of increasing concern for downstream analyses. To ensure that researchers have confidence in the data quality, we will continue to share the informative metrics computed by the standardized PDXNet quality control workflows.

Several key features are the focus for the next iteration of the PDXNet Portal. These include tools to search for commonly found genomic variants (ex. SNPs, INDELs, and copy number variations) within models, diseases and genes of interest, and interactive exploration of gene expression data. These tools would enable researchers to perform meaningful analyses directly from the portal and more rapidly realize value from PDXNet data. Visualization and analysis of associated data, including imaging and tumor volume/drug response data, will also be a focus. These data types have the potential for high impact, particularly given the innovation in large scale data visualization techniques in many fields.

Further development of links between the PDXNet Portal and NCI computational infrastructure will benefit researchers as well. Moving large quantities of data is time-consuming and can be expensive. Enabling researchers to perform their analysis where the data is already present lowers the entry barrier into the computational analysis of PDX model data. To facilitate these links, we envision users will be able to create cohorts for analysis using the PDXNet Portal and transferring that selection to the Cancer Genomics Cloud or other computational platform, where they will be able to easily take advantage of well-developed computational infrastructure.

We will extend PDXNet Portal capabilities as the size and complexity of PDXNet datasets grow. These enhancements will allow the research community to quickly find and evaluate PDXNet resources to supplement their research studies. We will continue to improve the PDXNet Portal value by collaborating with related PDX initiatives, including the PDMR, PDXFinder, and EuroPDX. Such collaborations will demonstrate how to effectively conduct studies across institutions, providing examples for the broader research community in how to optimize their PDX studies with respect to the public PDX models and datasets that are becoming increasingly available.

## Acknowledgments

We would like to thank the patients who provided the tissues that support the PDXNet model and sequencing data generation.

This work was supported by NIH funding to the PDXNet Data Commons and Coordination Center (NCI U24-CA224067) to the PDX Development and Trial Centers (NCI U54-CA224083, NCI U54-CA224070, NCI U54-CA224065, NCI U54-CA224076, NCI U54-CA233223, and NCI U54-CA233306). The Cancer Genomics Cloud powered by Seven Bridges is a Cancer Research Data Commons Cloud Resource, funded in whole or in part with Federal funds from the National Cancer Institute, National Institutes of Health, Contract No. HHSN261201400008C and ID/IQ Agreement No. 17X146 under Contract No. HHSN261201500003I and 75N91019D00024.

## Supplementary Materials

### PDXNet Member Contribution

#### PDXNet Leadership and Data Submission

AW, BDD, BW, CJB, CXP, DAD, FMB, JD, JM, JHC, JR, GW, LCC, LD, MD, MH, MSC, MTW, NM, PNR, SK, SL, TW, YAE

#### Data Science, Management and Processing

AS, BJS, BW, CF, SK, MWL, SS, DAD, JG, JHC, SLS, SN, PDXNet Consortium, YAE, XYW

#### Portal Development

AS, CF, DAD, JG, JHC, MWL, SK

#### Portal Integration Planning

SK, MWL, DAD, JG, MR, SLS, SS, JHC

#### Writing Manuscript

SK, MWL, DAD, JG, JHC

## Supplementary Tables

**Supplementary Table 1.** Metadata associated with Hematoxylin and eosin (H&E) Images on the PDXNet Portal.

| Hematoxylin and Eosin (H&E) Image Metadata |  |
| --- | --- |
| Age | Percent stromal content |
| Cell annotation available? | Percent tumor content |
| Contributor | Primary cancer site |
| CTEP Code | Proteomics |
| Diagnosis Subtype | Race |
| Disease Type | Regional annotation available? |
| Engraftment site | RNA-Seq/ Exp array |
| Ethnicity | Sample ID |
| Gender | Sample Type |
| Histology | SNP array |
| Image file name | Stain |
| Is information of this model already in CGC? | Staining/scanning method available? |
| Magnification | Thumbnail |
| Metastatic site | Treatment |
| Model ID | Treatment information in patient tumor |
| Mouse strain | Treatment information in PDX tumor |
| Note | Tumor Biomarkers |
| Original Species | Tumor differentiation |
| Other pathology notes | Tumor Grade |
| Passage | Tumor Stage |
| Patient ID | WES/Mutations |
| Percent necrotic content |  |

**Supplementary Table 2.**
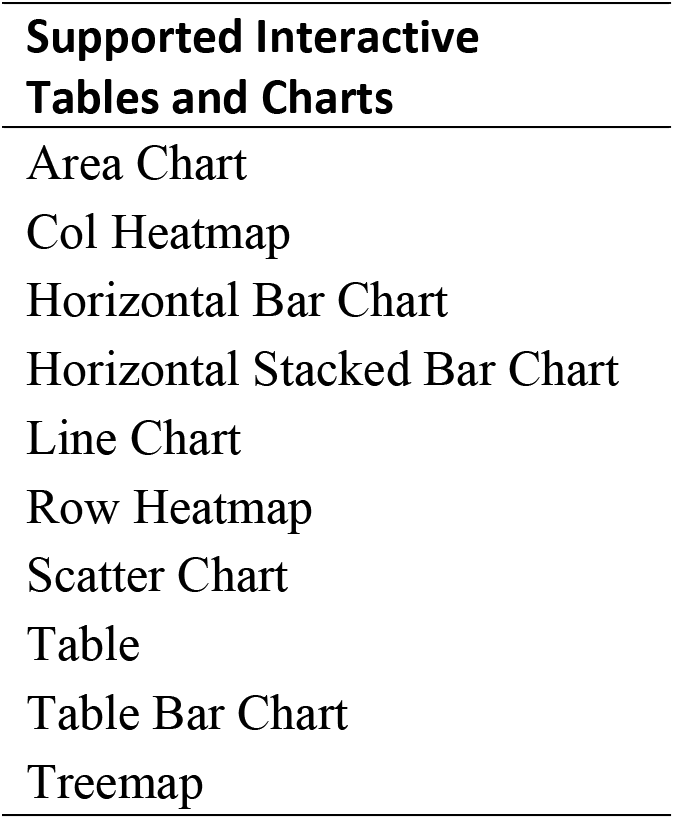
Supportive interactive table options on the PDXNet portal.

**Supplementary Table 3.**
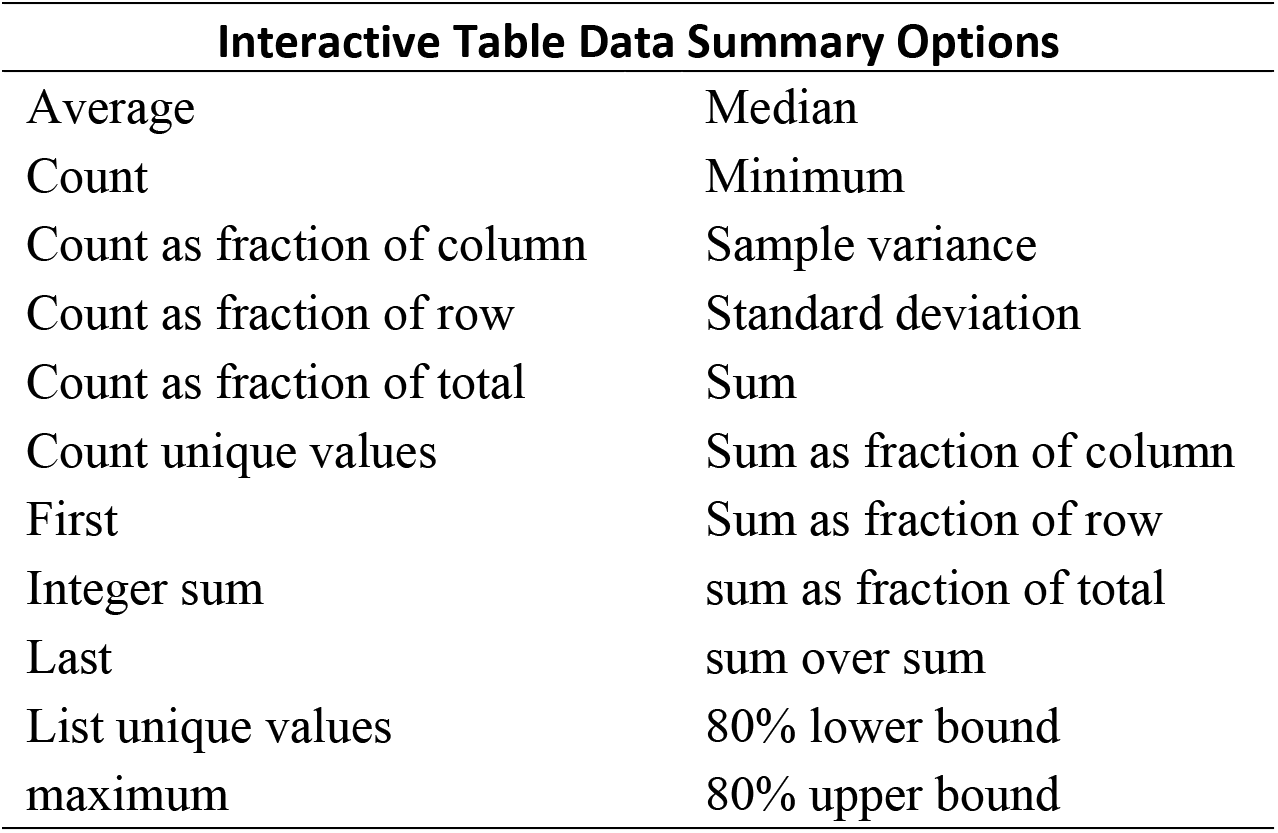
Data summary options available through interactive tables on the PDXNet Portal.

**Supplementary Table 4.** PDXNet metadata fields available on the Interactive Table Explorer on the PDXNet Portal.

| **PDXNet Metadata Fields Available on the**<br>**Interactive Table Explorer** |  |
| --- | --- |
| Access Level | File size |
| Availability | Gender |
| Capture Assembly | Investigations |
| Capture Kit | Is FFPE |
| Case id | Model id |
| Comments | Paired End |
| Contributor | Patient id |
| Created Date and time | Platform |
| Data Category | Public |
| Data Format | Sample ID |
| Data Type | Sample type |
| experimental strategy | Tumor id |
| File name |  |

**Supplementary Table 5.** Patient-derived model repository sequencing data processed with standardized PDXNet workflows referenced on the PDXNet Portal.

|  | **Overall** | **RNA-Seq** | **WES** |
| --- | --- | --- | --- |
| **Sample Type, n(%)** | N=9492 | N=4480 | N=5012 |
| Normal | 520(5.5%) | 0(0%) | 520(10%) |
| Organoid | 304(3.2%) | 152(3.4%) | 152(3.0%) |
| PDC | 276(2.8%) | 124(2.8%) | 152(3.0) |
| PDX | 7846(83%) | 3930(88%) | 3916(78%) |
| Tumor | 542(5.7) | 274(6.1%) | 268(5.3%) |
| unknown | 4(<0.1%) | 0(0%) | 4(<0.1%) |

**Supplementary Table 6.**
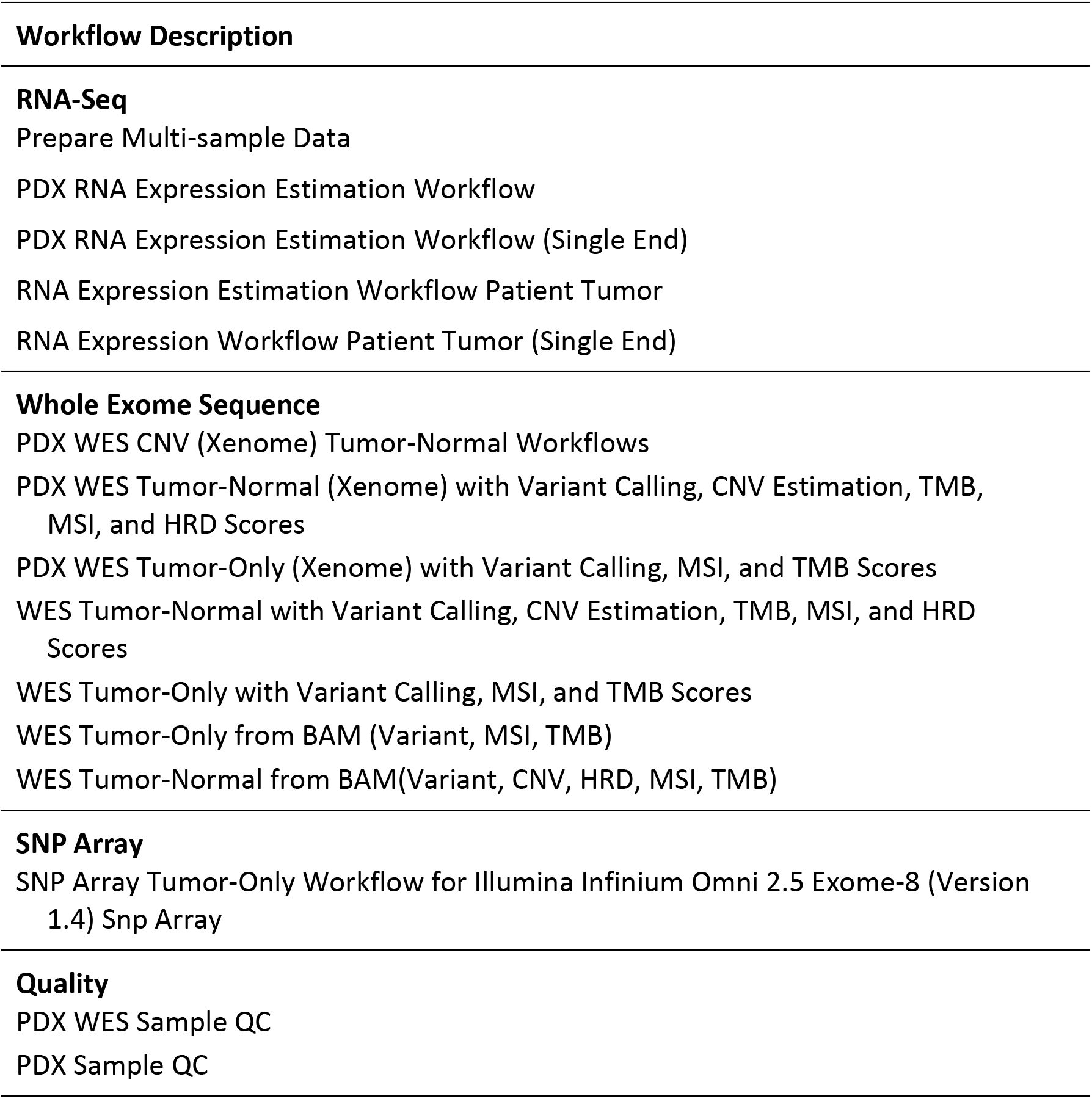
Standardized PDXNet bioinformatics workflows linked to the PDXNet Portal.

**Supplementary Table 7.** Summary of computed ancestry for PDXNet Models.

|  | Contributors |  |  |  | # of |
| --- | --- | --- | --- | --- | --- |
|  | HCI-BCM | MDACC | PDMR | WUSTL | Patients |
| AFR | 10 | 3 | 8 | 3 | 24 |
| Adenocarcinoma - colon |  |  | 4 |  | 4 |
| Adenocarcinoma - pancreas |  |  |  | 2 | 2 |
| Adenocarcinoma - rectum |  |  | 1 |  | 1 |
| Invasive breast carcinoma | 10 |  |  |  | 10 |
| Lung Adenocarcinoma |  | 3 | 2 |  | 5 |
| Melanoma |  |  | 1 | 1 | 2 |
| AMR | 18 | 2 |  |  | 20 |
| Invasive breast carcinoma | 18 |  |  |  | 18 |
| Lung Adenocarcinoma |  | 2 |  |  | 2 |
| EAS |  |  | 1 |  | 1 |
| Adenocarcinoma - colon |  |  | 1 |  | 1 |
| EUR | 8 | 30 | 158 | 24 | 220 |
| Adenocarcinoma - colon |  |  | 72 | 7 | 79 |
| Adenocarcinoma - pancreas |  |  | 24 | 13 | 37 |
| Adenocarcinoma - rectum |  |  | 14 | 1 | 15 |
| Invasive breast carcinoma | 8 |  | 10 | 2 | 20 |
| Lung Adenocarcinoma |  | 30 | 14 | 1 | 45 |
| Melanoma |  |  | 24 |  | 24 |
| MIX | 1 | 2 | 3 |  | 6 |
| Adenocarcinoma - colon |  |  | 1 |  | 1 |
| Adenocarcinoma - pancreas |  |  | 1 |  | 1 |
| Invasive breast carcinoma | 1 |  |  |  | 1 |
| Lung Adenocarcinoma |  | 2 | 1 |  | 3 |
| **# of Patients** | **37** | **37** | **170** | **27** | **271** |

HCI-BCM: Huntsman Cancer Center, MDACC: MD Anderson Cancer Center, PDMR: Patient-Derived Model Repository, WUSTL: Washington University at St. Louis

**Supplementary Table 8.** Summary of PDMR tumor volume dataset on the PDXNet portal shown by disease type and treatment.

| Disease | Treatment |  |  |  |  |  |  |  | Total |
| --- | --- | --- | --- | --- | --- | --- | --- | --- | --- |
|  | Control | 5-FU | Carboplatin | Erlotinib | Gemcitabine | Irinotecan | Paclitaxel | Vemurafenib |  |
| Colon | 13 | 10 | 10 | 7 | 11 | 11 | 11 | 7 | 81 |
| Lung | 13 | 9 | 10 | 8 | 10 | 10 | 10 | 8 | 78 |
| Pancreas | 12 | 11 | 1 | 9 | 11 | 11 | 11 | 7 | 82 |
| Skin | 9 | 7 | 1 | 3 | 7 | 6 | 7 | 3 | 48 |
| Bladder | 7 | 5 | 1 | 3 | 5 | 6 | 6 | 4 | 42 |
| Head and Neck | 10 | 5 | 1 | 4 | 6 | 6 | 6 | 3 | 45 |
| Kidney | 2 | 2 |  | 1 | 2 | 2 | 2 | 1 | 14 |
| Ovarian | 1 | 1 | 1 |  | 1 | 1 | 1 |  | 6 |
| Uterine | 4 | 4 | 1 | 1 | 4 | 4 | 4 | 1 | 26 |
| Gastric | 1 | 1 | 1 | 1 | 1 | 1 | 1 | 1 | 8 |
| Soft Tissue Neoplasm | 9 | 8 | 3 | 2 | 9 | 9 | 9 | 1 | 54 |
| Endocrine | 1 | 1 | 3 | 1 | 1 | 1 | 1 |  | 7 |
| Bone | 1 | 1 | 1 |  | 1 |  | 1 |  | 5 |
| CNS | 3 | 3 | 1 | 2 | 3 | 3 | 3 | 2 | 22 |
| Rectum | 1 | 1 | 1 |  | 1 | 1 | 1 |  | 6 |
| Grand Total | 87 | 69 | 36 | 42 | 73 | 72 | 74 | 38 | 524 |

## Graphical User Interface Screenshots

### Resources

**Supplement Figure 1.**
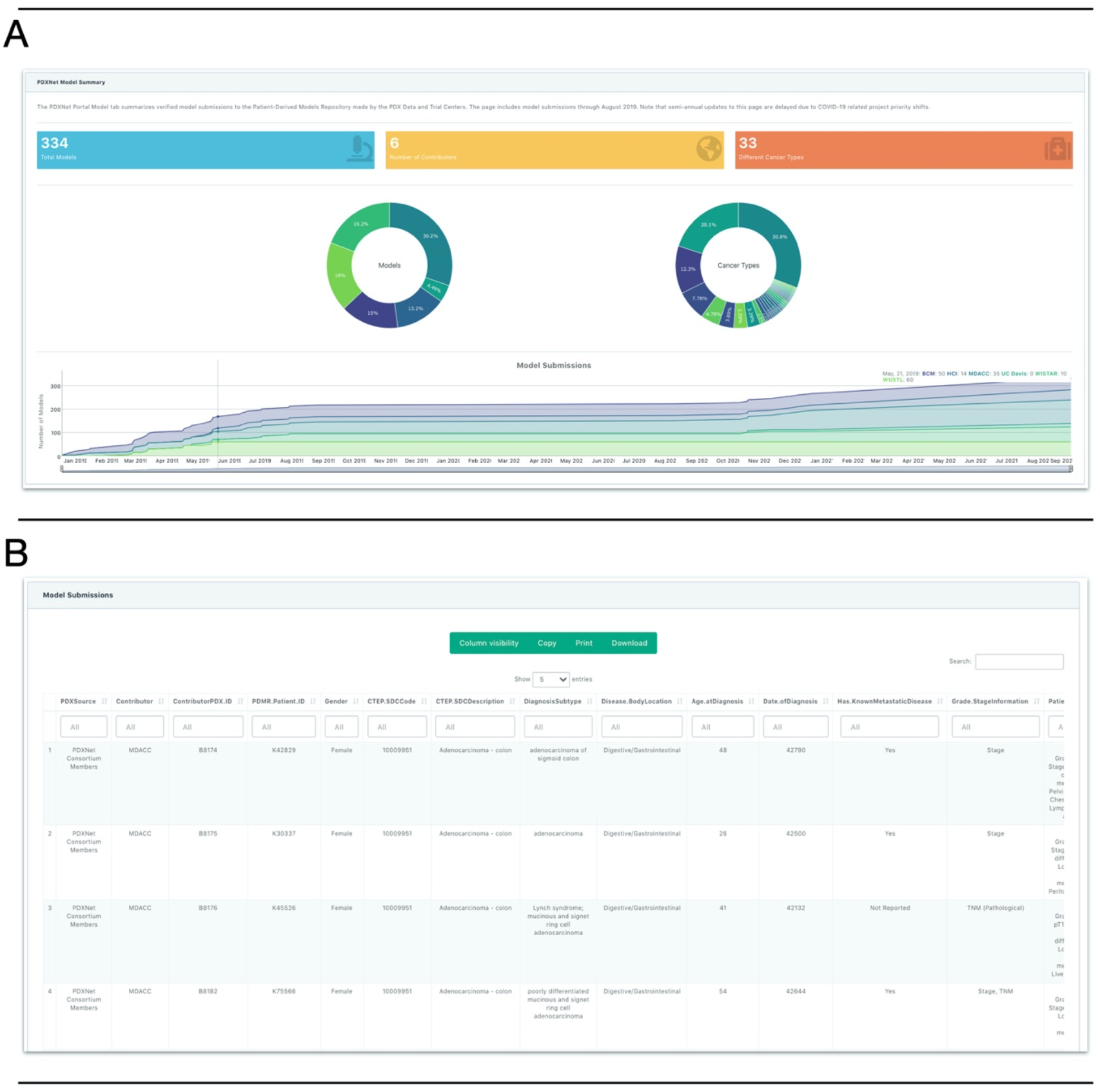
PDX models generated by PDXNet researchers shown on the PDXNet Portal. Figure shows components of the PDXNet Model sequencing data page in separate panels **(A)** Panel shows summary statistics including number of total models (blue), number of contributors (yellow), and number of cancer types (orange). Also, shown are donut plots for contributors and cancer types. Below the donut plots is a chart showing the number of models generated since January 2019. **(B)** Panel shows metadata for the PDXNet PDX models in a spreadsheet format. The interface supports searching and sorting metadata. Users can copy, print, and download metadata into accessible formats.

**Supplement Figure 2.**
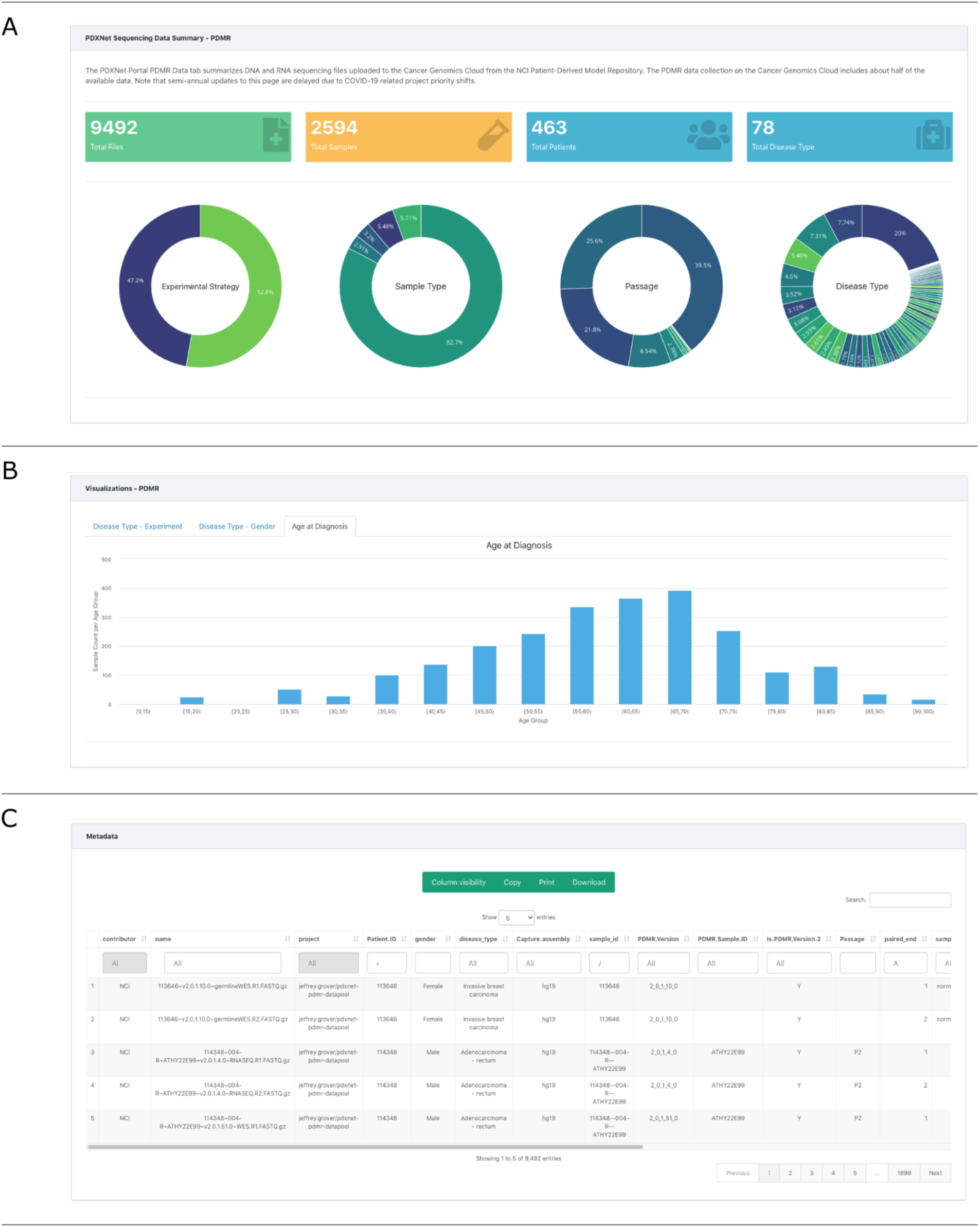
Patient Derived Model Repository (PDMR) sequencing data listed on the PDXNet portal. Figure shows components of the PDMR sequencing data page in separate panels **(A)** Panel shows summary statistics including number of sequencing files (green), contributors (yellow), total samples (orange), and total patients (blue). Also, shown are donut plots for contributors, sample types, disease type, experimental strategy, WES contributors, and RNA-Seq contributors. **(B)** Panel shows age from PDMR patients on a bar chart. **(C)** Panel shows metadata for the PDMR sequencing data in a spreadsheet format. The interface supports searching and sorting metadata. Users can copy, print, and download metadata into accessible formats.

**Supplement Figure 3.**
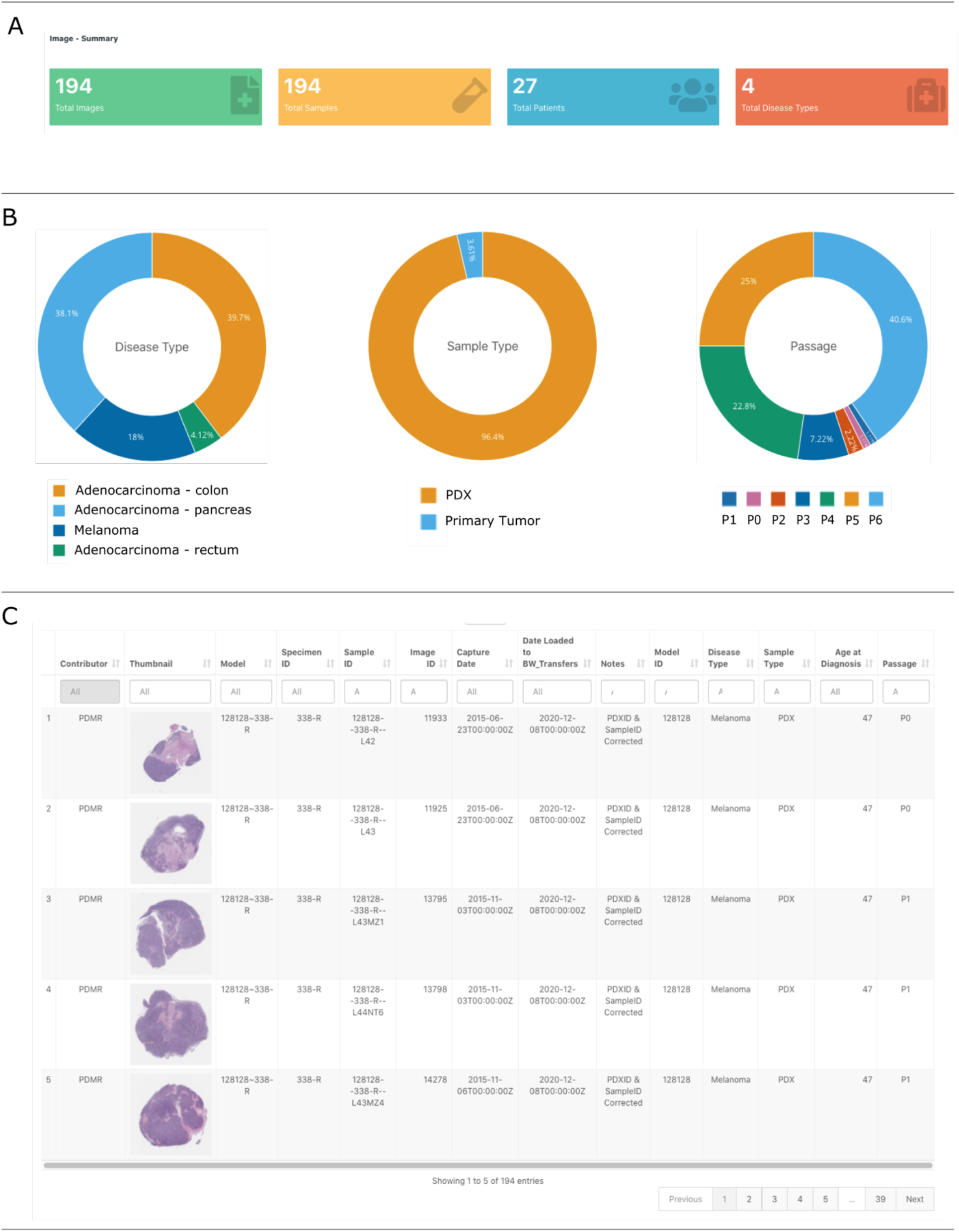
Patient-Derived Model Repository (PDMR) image page on the PDXNet Portal. Figure shows components of the PDMR image data page in separate panels **(A)** Panel shows summary statistics including number of images (green), contributors (yellow), total patients (blue), and total disease types (red). **(B)** Panel B shows donut plots for disease type, sample type, and passage. **(C)** Panel shows metadata for the PDMR image data in a spreadsheet format. The interface supports searching and sorting metadata. Users can copy, print, and download metadata into accessible formats.

**Supplement Figure 4.**
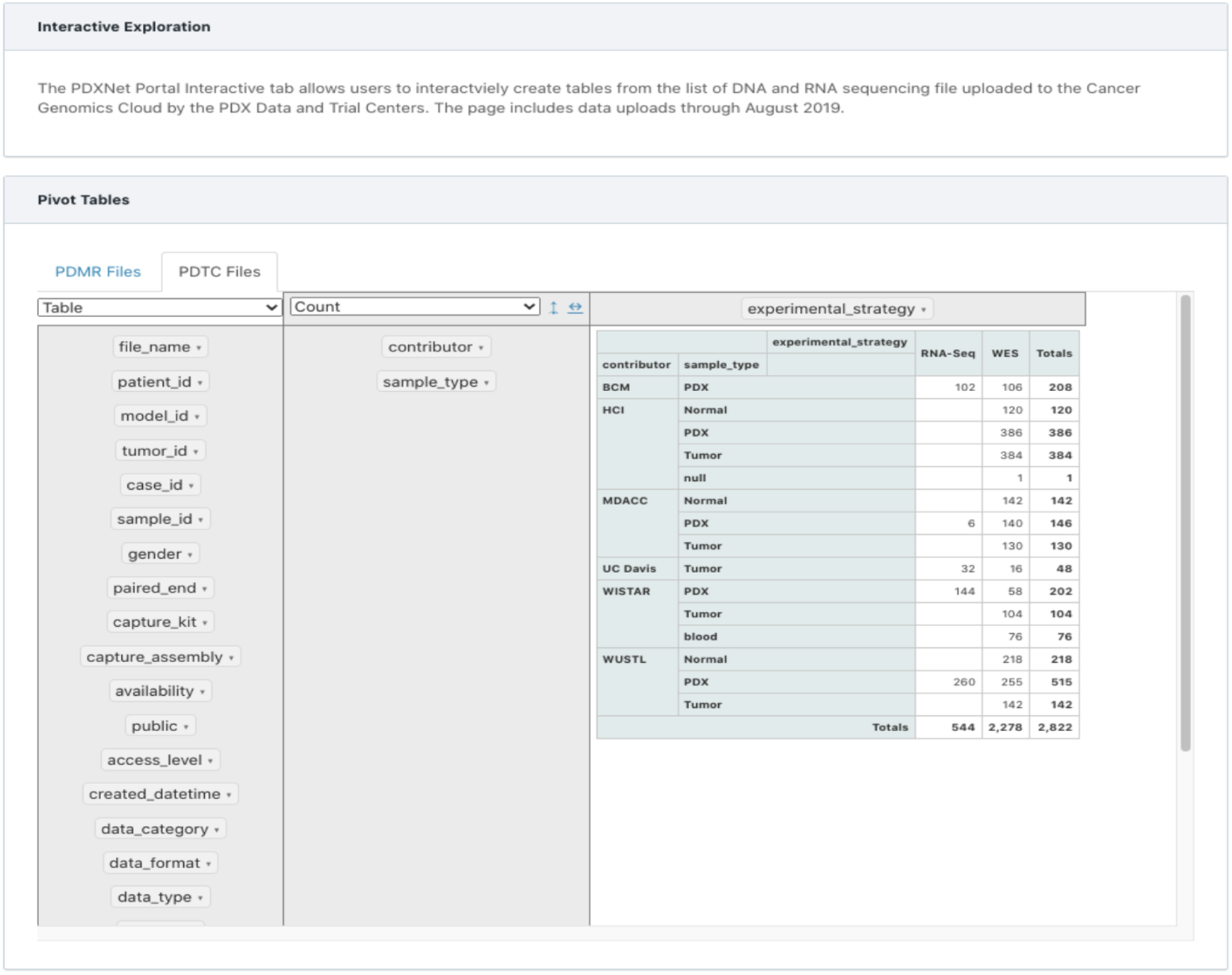
Interactive data explorer page on the PDXNet Portal. Figure shows the interactive exploration page on the PDXNet portal. The user can interactively create a pivot table with either the metadata from the PDXNet sequencing data or the PDMR sequencing data. Constructing the table involves dragging and dropping table fields on the left side to the table area (green) on the right side of the screen.

**Supplement Figure 5.**
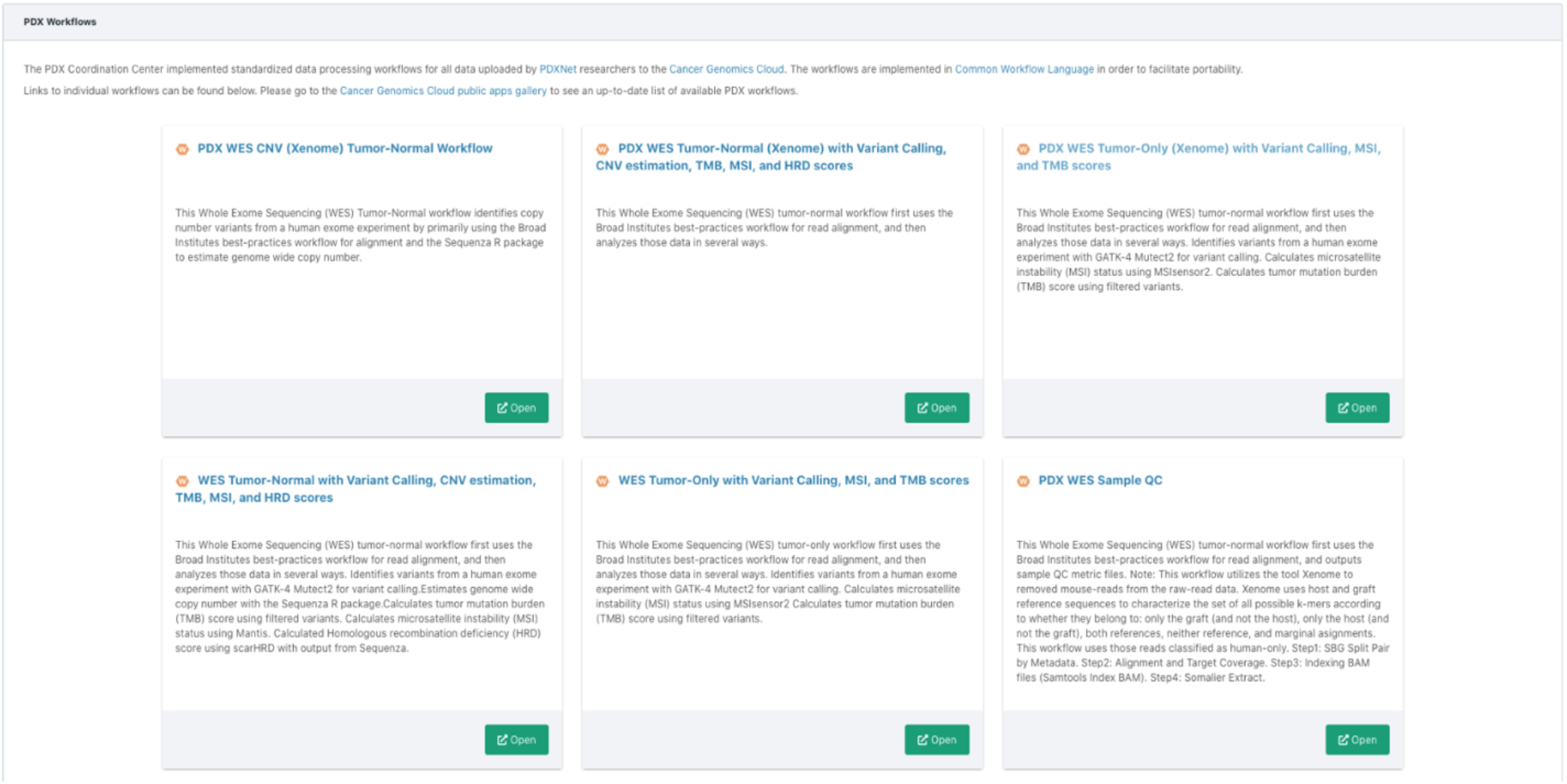
Standardized PDXNet processing workflows shown on the PDXNet Portal. Figure shows a section of the workflow page on the PDXNet portal. The page includes brief descriptions of standardized workflows created to process RNA-Seq, whole exome, and to a lesser extent array data. The page includes links to comprehensive workflow documentation on the Cancer Genomics Cloud Public Apps Gallery; where the workflows are made publicly available.

**Supplement Figure 6.**
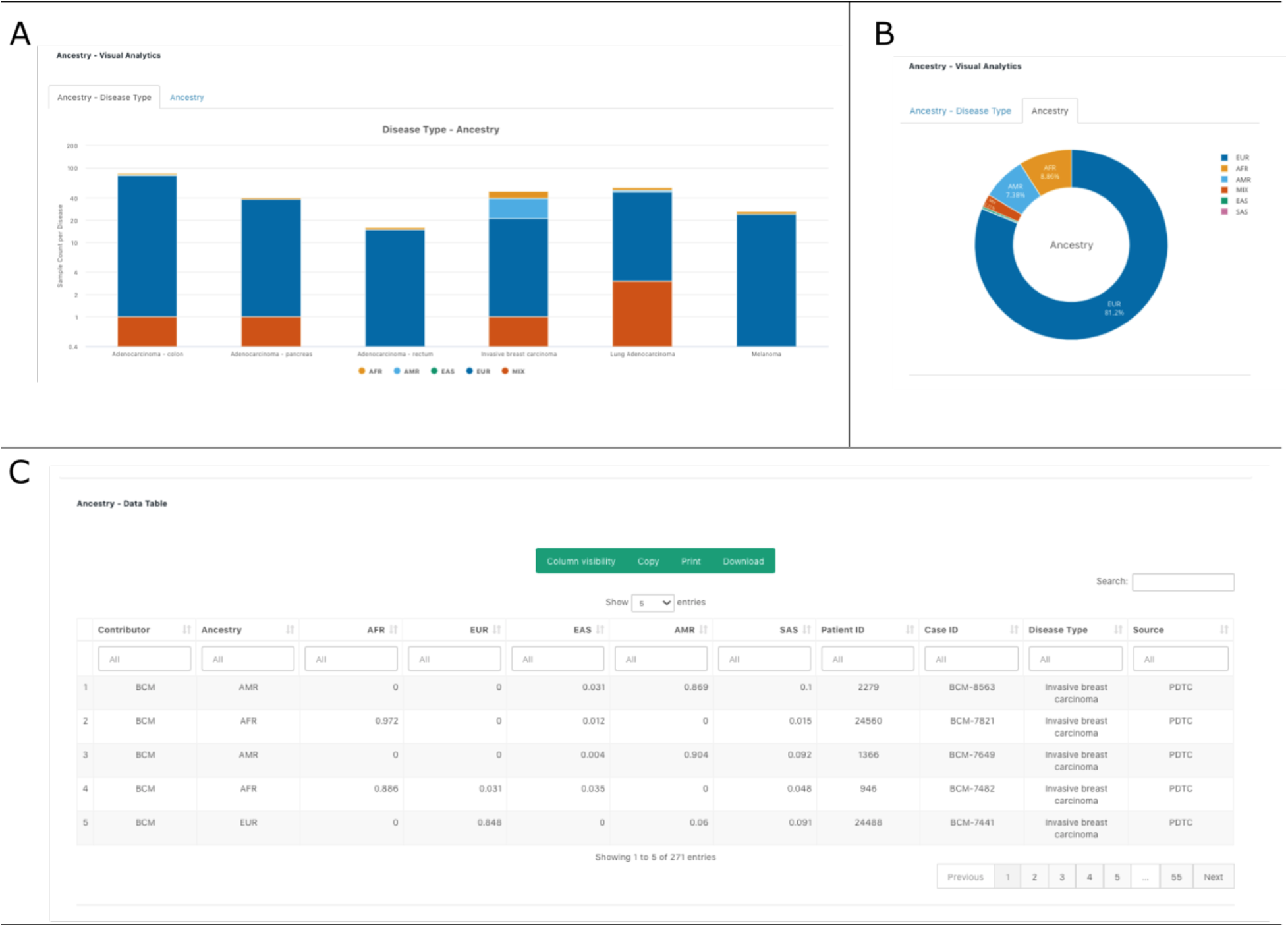
Ancestry information computed from sequencing data shown on the PDXNet Portal. Figure shows plots generated on the ancestry data page of the PDXNet portal **(A)** Panel A shows a stacked bar chart with each bar corresponding to a user selected disease. Each bar shows the ancestry composition of available samples by color. The ancestry algorithm classifies samples as African (AFR), American (AMR), East Asian (EAS), South Asian (SAS), and Mixed (MIX). **(B)** Panel shows computed ancestry in a pie chart. Ancestry is classified in the following categories: European(EUR), African(AFR), American(AMR), Mixed(MIX), East Asian(EAS), South Asian(SAS). **(C)** Panel shows ancestry metadata for the processed sequencing data in a spreadsheet format. The interface supports searching and sorting metadata. Users can copy, print, and download metadata into accessible formats.

**Supplement Figure 7.**
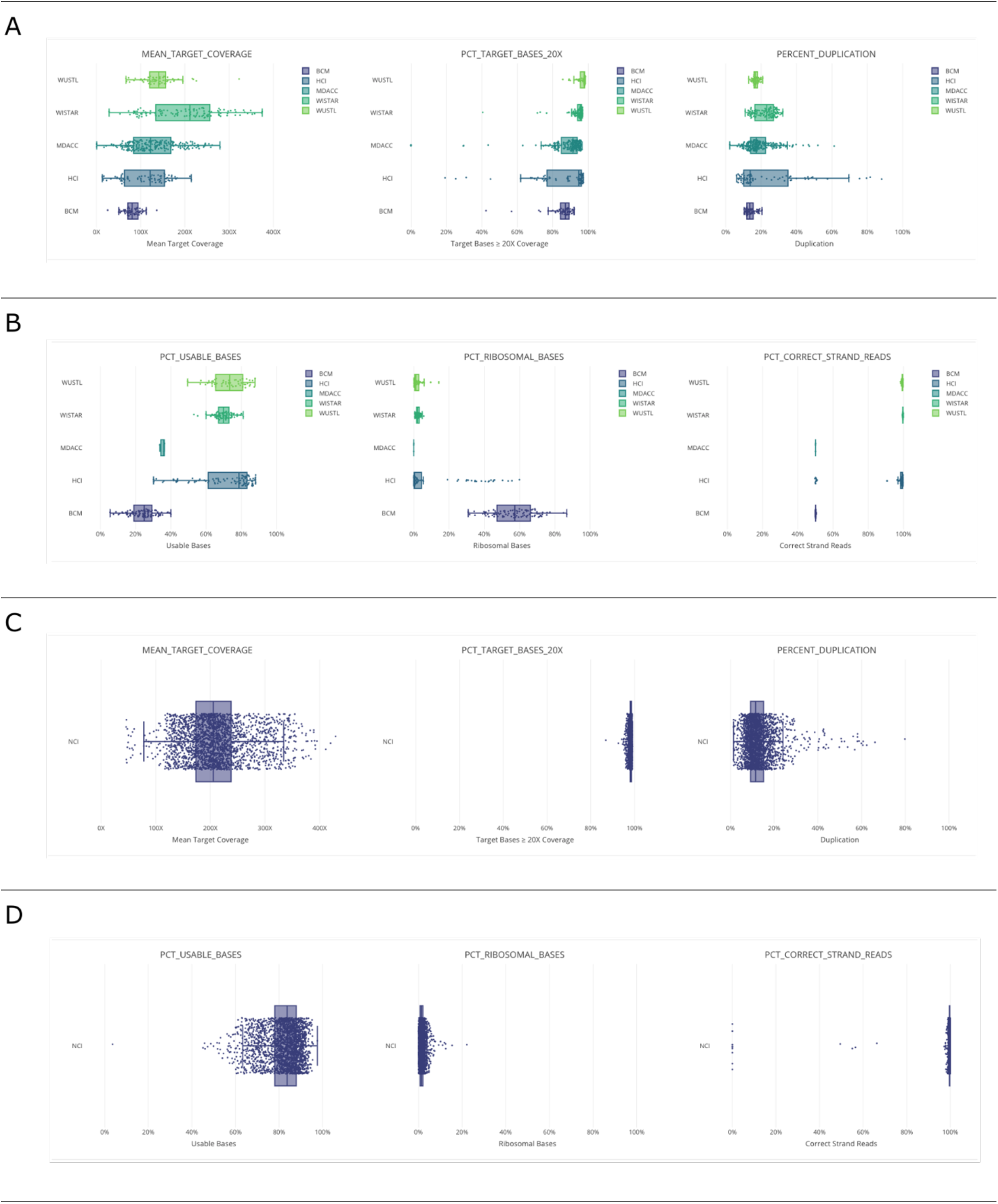
Sequencing quality control plots examples generated on the PDXNet Portal. Figure shows sequencing data QC figures generated on the quality control page of the PDXNet Portal (A) Plot shows mean target coverage, percent target coverage at 20x, and percent duplication as box plots with data for each PDX Development and Trial Center presented as a different box plot. (B) Plot shows percent usable basis, percent ribosomal basis, and percent correct strand reads as box plots with data for each PDX Development and Trial Center presented as a different box plot. (C) Plot shows mean target coverage, percent target coverage at 20x, and percent duplication as box plots generated from PDMR data. (D) Plot shows percent usable basis, percent ribosomal basis, and percent correct strand reads as box plots generated from PDMR data.

## References

1. Koga, Y. & Ochiai, A. Systematic Review of Patient-Derived Xenograft Models for Preclinical Studies of Anti-Cancer Drugs in Solid Tumors. Cells 8, 418 (2019).

2. Dobrolecki, L., Airhart, S., Alferez, D. & Aparicio, S. Patient-derived Xenograft (PDX) models in basic and translational breast cancer research. Cancer Metastasis Rev. 35, 547–573 (2016).

3. Collins, A. T. & Lang, S. H. A systematic review of the validity of patient derived xenograft (PDX) models: The implications for translational research and personalised medicine. PeerJ 2018, 1–22 (2018).

4. Brown, K. M. et al. Patient-derived xenograft models of colorectal cancer in preclinical research: A systematic review. Oncotarget 7, 66212–66225 (2016).

5. Grandori, C. & Kemp, C. Personalized cancer moels for target discovery and precision medicine. Trends Cancer 4, 634–642 (2018).

6. Hidalgo, M. et al. Patient-Derived Xenograft Models: An Emerging Platform for Translational Cancer Research. Cancer Discovery 4, 998–1013 (2014).

7. Woo, X. Y. et al. Genomic data analysis workflows for tumors from patient-derived xenografts (PDXs): Challenges and guidelines. BMC Medical Genomics 12, 1–19 (2019).

8. Evrard, Y. A. et al. Systematic establishment of robustness and standards in patient-derived xenograft experiments and analysis. Cancer Research 80, 2286–2297 (2020).

9. Sun, H. et al. Comprehensive characterization of 536 patient-derived xenograft models prioritizes candidates for targeted treatment. Nature Communications 12, (2021).

10. Woo, X. Y. et al. Conservation of copy number profiles during engraftment and passaging of patient-derived cancer xenografts. Nature Genetics 53, 86–99 (2021).

11. The NCI Patient-Derived Models Repository (PDMR). Research, NCI-Frederick: Frederick National Laboratory for Cancer https://pdmr.cancer.gov/ (2021).

12. Conte, N. et al. PDX Finder: A portal for patient-derived tumor xenograft model discovery. Nucleic Acids Research 47, D1073–D1079 (2019).

13. Meehan, T. F. et al. PDX-MI: Minimal information for patient-derived tumor xenograft models. Cancer Research 77, e62–e66 (2017).

14. Lowy, D. & Collins, F. Aiming High - Changing the Trajectory for Cancer. NEJM 374, 1901–1904 (2016).

15. Sharpless, N. E. & Singer, D. S. Progress and potential: the Cancer Moonshot. Cancer Cell 1–6 (2021) doi:10.1016/j.ccell.2021.04.015.

16. Lau, J. W. et al. The cancer genomics cloud: Collaborative, reproducible, and democratized - A new paradigm in large-scale computational research. Cancer Research 77, e3–e6 (2017).

17. NCI. Cancer Data Service. https://datacommons.cancer.gov/repository/cancer-data-service (2021).

18. Amstutz, P., Crusoe, M. & Tijianic, N. Common Workflow lanaguage (CWL) Workflow Description, v1.0.2. (2016).

19. Chen, C. Y. et al. Improved ancestry inference using weights from external reference panels. Bioinformatics 29, 1399–1406 (2013).

20. Auton, A. et al. A global reference for human genetic variation. Nature 526, 68–74 (2015).

21. Weiss, K. M. & Long, J. C. Non-Darwinian estimation: My ancestors, my genes’ ancestors. Genome Research 19, 703–710 (2009).

22. Sztupinszki, Z. et al. Migrating the SNP array-based homologous recombination deficiency measures to next generation sequencing data of breast cancer. npj Breast Cancer 4, 8–11 (2018).

23. Cingolani, P. et al. A program for annotating and predicting the effects of single nucleotide polymorphisms, SnpEff: SNPs in the genome of Drosophila melanogaster strain w1118; iso-2; iso-3. Fly 6, 80–92 (2012).

24. Kautto, E. A. et al. Performance evaluation for rapid detection of pan-cancer microsatellite instability with MANTIS. Oncotarget 8, 7452–7463 (2017).

25. Niu, B. et al. MSIsensor: Microsatellite instability detection using paired tumor-normal sequence data. Bioinformatics 30, 1015–1016 (2014).

26. Hastie, T., Tibshirani, R. & Friedman, J. Boosting and Additive Trees. in The Elements of Statistical Learning (Springer, 2009).

27. Mailman, M. D. et al. The NCBI dbGaP database of genotypes and phenotypes. Nature Genetics 39, 1181–1186 (2007).

28. Lau, J. W. et al. The cancer genomics cloud: Collaborative, reproducible, and democratized - A new paradigm in large-scale computational research. Cancer Research 77, e3–e6 (2017).

